# Comparative phylotranscriptomics reveals a 110 million years-old symbiotic program

**DOI:** 10.1101/2022.09.02.505815

**Authors:** Cyril Libourel, Jean Keller, Lukas Brichet, Anne-Claire Cazalé, Sébastien Carrère, Tatiana Vernié, Jean-Malo Couzigou, Caroline Callot, Isabelle Dufau, Stéphane Cauet, William Marande, Tabatha Bulach, Amandine Suin, Catherine Masson-Boivin, Philippe Remigi, Pierre-Marc Delaux, Delphine Capela

## Abstract

Symbiotic interactions have structured past and present ecosystems and shaped the evolution of life. As any trait, the symbiotic state observed in extant species builds on ancestral and conserved features, and lineage-specific innovations. From these mixed origins, defining the ancestral state of symbiotic associations is challenging although it is instrumental for understanding how symbiotic abilities emerge from non-symbiotic states. Here we aimed at reconstructing the intermediate steps leading to the root-nodule nitrogen-fixing symbiosis (RNS) observed in some extant flowering plants. For this, we compared the transcriptomic responses of nine host plants in response to symbiotic bacteria. We included the mimosoid legume *Mimosa pudica* for which we assembled a chromosome-level genome and generated the transcriptomic response to experimentally evolved bacterial symbionts. With this dataset, we reconstructed the ancestral RNS transcriptome, composed of most already described symbiotic genes together with hundreds of novel candidates. We found that the response to the chemical signals produced by the symbiont, nodule organogenesis and nitrogen-fixation are predominantly linked to ancestral responses, although these traits have diversified in the different nitrogen-fixing lineages. We detected a clear signature of recent and convergent evolution for the ability to release intracellular symbiosomes in two legume lineages, exemplified by the expression of different classes of small proteins in each group, potentially leading to the convergent gain of symbiotic evolutionary stability. Our findings demonstrate that most of the novelties for RNS were mostly in place in the most recent common ancestor of the RNS-forming species that lived on Earth 110 million years ago.

**Graphical abstract:** A little graphical/nice phylogeny with nodes of interest

**Highlights:**

- We sequenced a high-quality genome of the Mimosoideae *Mimosa pudica*
- The nitrogen-fixing root-nodule symbiosis relies on an ancestral transcriptomic response
- All symbiotic traits involve genes of the ancestral symbiotic program
- Symbiont perception, nodule organogenesis and nitrogen-fixation are essentially ancestral processes
- Convergent evolution of intracellular accommodation of symbionts additionally involves lineage-specific genes

## Introduction

Interactions between organisms forms a continuum of associations, from parasitism to mutually beneficial symbioses^1^, which have contributed for billions of years to the evolution and diversification of the plant lineage^2^. The mutualistic symbioses formed with fungal or bacterial symbionts are associated with key transitions, such as the colonization of lands by plants 450 million years ago which was enabled by the evolution of the arbuscular mycorrhizal symbiosis^3,4^. Following this initial event, plants and their symbiotic abilities have diversified leading to the emergence of multiple types of mutualistic symbioses with microorganisms^2^, which can be divided in two main groups, the intracellular and the extracellular symbioses. Extracellular symbioses include for instance plant – cyanobacteria interactions where the bacterial symbiont is hosted in dedicated canals and glands^5,6^, ectomycorrhizal symbioses between plant roots and ascomycetes or basidiomycetes fungi^7^, or very specific association between *Dioscorea sansibarensis* and its bacterial symbiont restricted to leave drip tips^8^. Intracellular symbioses are almost exclusively formed between plants and fungal symbionts^9^, with the nitrogen-fixing root nodule symbiosis (RNS) being the almost unique case of intracellular accommodation of bacteria^9^. However, intracellular accommodation of symbionts during RNS refers to different physical structures depending on the plant species, including bacteria being either retained in transcellular tubular structures^10^ or completely released in host cells forming organelle-like structures called symbiosomes^11^.

In extant species, RNS is found in ~17 500 species from four orders of flowering plants^12^, ~17 300 sp. from the Fabales and 230 species from the Fagales, Cucurbitales and Rosales, which together form the Nitrogen-Fixing Nodulation (NFN) clade^13^. Comparative phylogenomic studies coupled with previous phylogenetic and physiological work have revealed the evolutionary history of RNS. The most likely scenario proposes that RNS was gained only once, before the diversification of the NFN clade. Following that single gain, RNS diversified in each lineage, and was lost subsequently multiple times leading to the scattered distribution observed in extant species^13–15^. The rate of RNS loss differs between lineages, with some displaying an evolutionary stabilized association while others seem to have experienced massive losses. However, the nature of the ancestral RNS, its functioning and how it diversified over 110 million years of evolution remains elusive.

The comparison of transcriptomic patterns across species in a given context, being developmental or in response to the environment, allows reconstructing ancestral and derived responses to that context. For instance, this has been used to reconstruct the flooding response in angiosperms^16^, the evolution of the shoot meristem in plants^17^, and to infer the ancestral arbuscular mycorrhizal transcriptome, another type of plant symbiosis^4^.

Here, we combined transcriptomics in multiple species and phylogenomics to reconstruct the ancestral RNS transcriptome. We further dissected the transcriptional shifts associated with each symbiotic step by exploiting bacterial mutants and strains arising from experimental evolution^18^, which progressively recapitulates the full symbiotic interaction. Together, our results provide new insights on the evolutionary origins of the different symbiotic stages in a complex mutualistic interaction.

## Results and Discussion

### Identification of an ancestral RNS transcriptomic signature

The two largest groups of RNS-forming species are the Mimosoideae and Papilionoideae sub-families in the Fabales order^15^. While transcriptomic data have been obtained in response to RNS in a number of Papilionoideae, the Mimosoideae have been ignored. To fill this gap, we conducted a time-course experiment with *Mimosa pudica* inoculated with its bacterial symbiont *Cupriavidus taiwanensis* (Table S1). In order to determine differentially expressed genes accurately, we *de novo* sequenced the genome of *M. pudica* using a combination of long-read sequencing and optical mapping (See Material and Methods) leading to a near-chromosome level assembly (Table S2). This method allowed us to generate 74 hybrid scaffolds (from 128 kbp to 25.5 Mbp with N50 = 16.1 Mbp), for a total genome size of 797.25 Mb. Automated structural annotation of the genome yielded 73,541 protein coding genes and 5,134 ncRNAs. Finally, the high completeness of the annotated genome was evidenced by a 97% (2255 genes) Busco recovery score on eudicots_odb10 (C:97.0% [S:10.2%,D:86.8%], F:1.1%,M:1.9%,n:2326). The expression of 51,214 *Mimosa* genes was detected in our complete transcriptomic dataset, 43% of which were differentially expressed (8303 genes up/15423 genes down) during the symbiotic interaction with *C. taiwanensis* in at least one time point compared to non-inoculated roots (Table S3 and S4).

In addition, we generated the transcriptome of the Papilionoideae *Lupinus albus*, from the Genisteae tribe, inoculated with *Bradyrhizobium sp.* 1AE200 strain (Ledermann and Couzigou, unpublished) which forms lupinoid nodules^19^ and whose genome has been recently sequenced ^20^. In brief, we identified 3794/4781 (up/down) deregulated genes in response to *Bradyrhizobium sp*. 1AE200 compared to non-inoculated roots (Table S1, S3 and S4). These differentially expressed genes represent around 33% of the 26,204 *Lupinus* expressed genes.

To obtain comparable datasets, raw RNAseq reads obtained in the presence or absence of their respective bacterial symbionts from 7 other nodulating species (Table S1) were remapped on their respective genomes and differentially expressed genes (DEGs) were computed following the same approach as for *M. pudica* and *L. albus*. Due to sampling and sequencing depth heterogeneity among species, we used different fold-change thresholds to obtain comparable number of differentially expressed genes (See Material and Methods). For each species, we also concatenated all differentially expressed genes at any time point to estimate the whole symbiotic response for up and down regulated genes.

Between 2258/8303 differentially up- and 1895/15423 down-regulated genes were detected in the nine sampled species at any time of the symbiotic interaction (Table S3, S4 and S6). The whole symbiotic response represents 13% to 28% of upregulated genes (mean=20%) and 11% to 31% downregulated genes (mean=22%), representing an important transcriptomic response for all of the species considered. As expected, species with transcriptomic responses for only mature nodule, such as *Hippophae rhamnoides*, *Datisca glomerata* and *Lupinus albus*, exhibited a lower proportion of deregulated genes (Table S3 and S6).

Although, we observed a massive symbiotic transcriptomic response in all species, this might reflect either a conserved or species-specific responses, or a mix of both patterns. To determine the evolutionary origin of these responses, we computed orthogroups^21^ for the nine studied species, together with 16 additional species from the NFN clade and *Arabidopsis thaliana* as outgroup (Table S5). The additional species were chosen based on genome quality and to cover RNS- and non-RNS-forming clades. By cross-referencing the transcriptomic data with the orthogroups, we identify for each orthogroups which species contains at least one gene up/down regulated. These DEG-containing orthogroups will be named DEOG thereafter (Differentially Expressed OrthoGroups). Considering the DEOGs, we were able to reconstruct ancestral states for a discrete trait (species containing the DEOG, or not) using a maximum parsimony approach^22–24^. Using this method, we determined at which phylogenetic node the genes became differentially regulated during evolution (See Material and Methods). Besides expression itself, the number of predicted DEOGs at a given node depends on a number of factors such as the maximum number of orthogroups present at that node or the accuracy of the orthogroup reconstruction method. To consider these biases, we assessed whether the experimentally determined values for each node significantly deviate from random expectation (See Material and Methods). For each species, most of the genes were found differentially regulated in a species-specific manner (Figure 1, Table S6). However, the observed numbers were either not significantly different from the null expectation, or lower than expected (Table S6). By contrast, a number of internal nodes displayed significantly more DEOGs than expected (p-value < 0.001, Table S6). In particular, a total of 771/1481 orthogroups (151%/167% increase compared to the mean null expectation) were inferred to have been already up/down-regulated in the most recent common ancestor of all RNS-forming species. Among the 771 ancestrally up-regulated orthogroups, 157 contain genes with a known function, such as the common symbiotic pathway members LjNFR5/MtNFP^25,26^, SYMRK/DMI2^27,28^, CCaMK/DMI3^29^, CYCLOPS/IPD3^30^, the master symbiotic regulator NIN^31^ or the infection-associated gene RPG^32,33^. The phylogenetic distribution of three of them, LjNFR5/MtNFP, NIN and RPG, has been recently linked with the ability to form RNS in the NFN clade. Indeed, all three genes have been lost independently in multiple lineages no longer able to form the RNS^14,15^. In addition to the known genes, 614 orthogroups with undescribed functions were detected, including 131 that are absent from at least one non-RNS-forming species and present in all RNS-forming species, and thus represent top candidates for future investigation by reverse genetics (Figure 1, Table S7).

**Figure 1:**
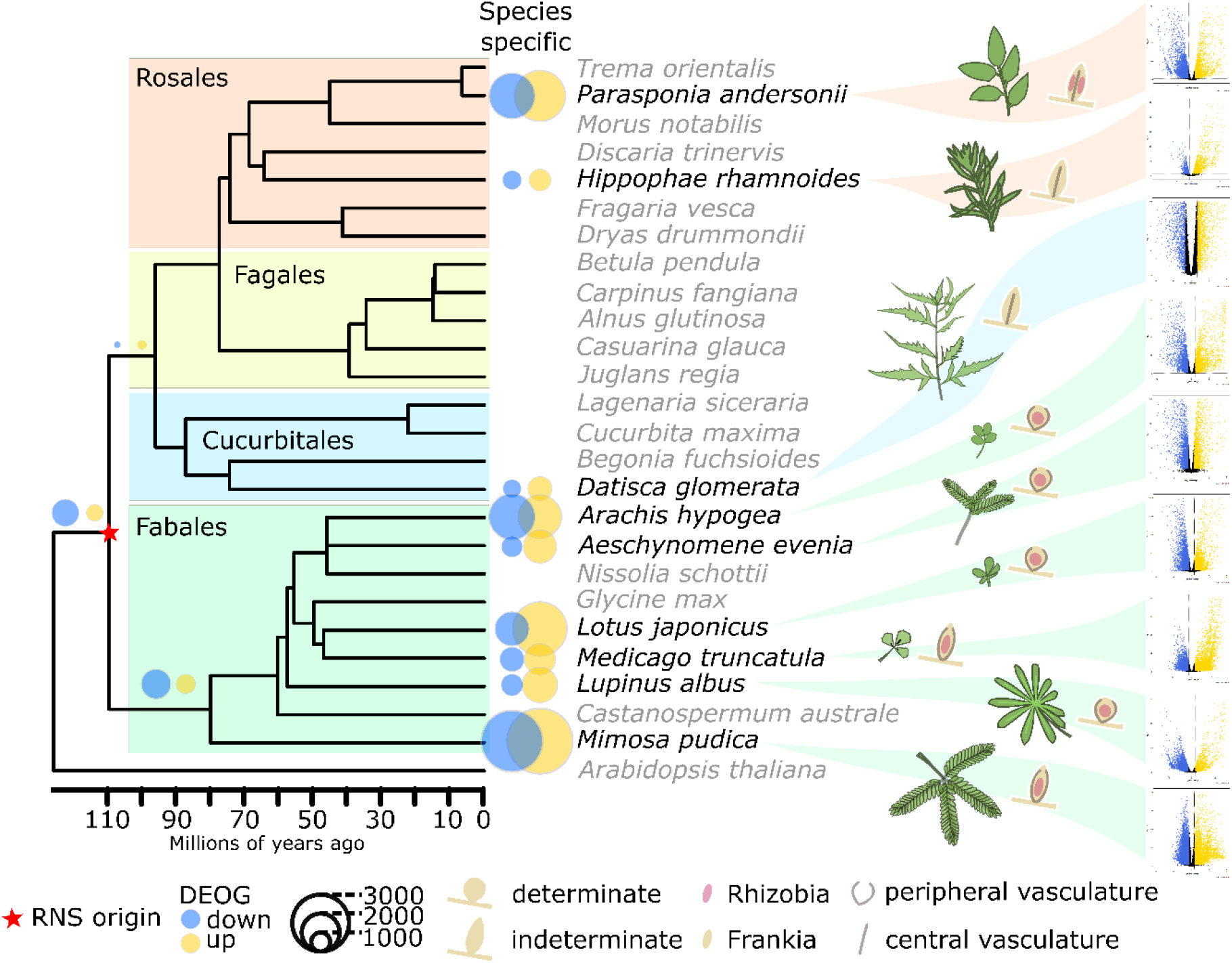
Conservation of the RNS transcriptomic response across NFN species. The tree depicts the orthofinder NFN phylogeny with *Arabidopsis thaliana* as outgroup. Species used to compare symbiotic transcriptomes are indicated in black, species used to compute orthogroups are indicated in gray. DEOG: Differentially Expressed genes containing OrthoGroups. The volcano plots on the right represent the logFC (x axis) by FDR *p*-values (y axis) for the 9 species at the latest time point. Blue and gold dots indicate significant down-regulated and up-regulated genes, respectively.

In other words, the combination of transcriptomic responses and phylogenomics allowed us to identify a set of 771 orthogroups that we can consider as the ancestral RNS transcriptome. We also identified orthogroups and genes recruited during the diversification of RNS in independent lineages, a pattern reminiscent of the potentiation – actualization – refinement model proposed for the evolution of novelties^34,35^.

### Gradual recruitment of plant symbiotic gene expression in experimentally evolved symbionts

RNS is a complex interaction involving multiple physiological and developmental processes, that are often coupled and overlapping. In the case of most Fabales, these processes include the perception and response to the symbiotic signal produced by the symbionts (the so-called Nod Factors, NF), nodule organogenesis and the concomitant penetration of bacteria within root and nodule tissues, the intracellular release and persistence of bacteria within plant cells, symbiosomes formation, and nitrogen fixation. The evolutionary transition from a non-RNS forming state to a fully functional RNS state likely occurred over millions of years through a number of intermediate stages which cannot be captured in extant species. To define the transcriptional modules (and their evolutionary origin) associated with each process, we exploited a collection of bacterial mutants that gradually induce the full symbiotic program. Most of these bacterial mutants originate from an evolution experiment that was developed to try mimicking the evolution of symbiotic abilities in a legume symbiont^36–38^. Starting from a non-symbiotic soil-borne bacterium, the *Ralstonia solanacearum* GMI1000 strain, in which we introduced the symbiotic plasmid pRalta (GMI1000+pRalta) from the *M. pudica* natural symbiont *C. taiwanensis* LMG19424^39^, we have propagated bacteria for 400 generations along successive nodulation cycles on *M. pudica.* All along the experiment, clones have gradually gained symbiotic abilities^36,38,40^ and adaptive mutations responsible for the main phenotypic changes observed in the evolved clones were identified. RNS was obtained following mutations inactivating the Type Three Secretion System (TTSS) of *R. solanacearum*. A stop mutation in *hrcV*, a gene encoding a TTSS structural protein, conferred to bacteria the capacity to nodulate *M. pudica* but nodules were only extracellularly invaded (Figure 2A). By contrast, a stop mutation in *hrpG*, a gene encoding a global regulator of hundreds of genes including TTSS genes, enabled bacteria to form nodules and invade them intracellularly through the formation of symbiosomes, which are released in the cytoplasm of nodule cells^36^. However, bacteria mutated in *hrpG* degenerate very rapidly following symbiosome release (Figure 2A). Cumulating a *hrpG* mutation with a mutation in the regulator *efpR* enhanced intracellular persistence of bacteria although to a level not yet equivalent to a wild-type or a non-fixing mutant of *C. taiwanensis*^38^ and was not yet sufficient to enable nitrogen-fixation in symbiosis with *M. pudica*. We reconstructed the adaptive mutations *hrcV*, *hrpG* and *hrpG-efpR* in the non-symbiotic original strain GMI1000+pRalta to generate a collection of nearly-isogenic strains with increased symbiotic abilities (Figure 2A). We analyzed the transcriptome of *M. pudica* in response to inoculation with each of these three mutants, the non-nodulating parental strains GMI1000, GMI1000+pRalta, and a *nifH* mutant of *C. taiwanensis,* which is only affected in its ability to fix nitrogen^41^. We harvested tissue samples, either roots, nodules primordia or nodules, at different time points between 1 and 21 days after inoculation to capture the most advanced symbiotic response induced by each mutant (Table S1).

**Figure 2:**
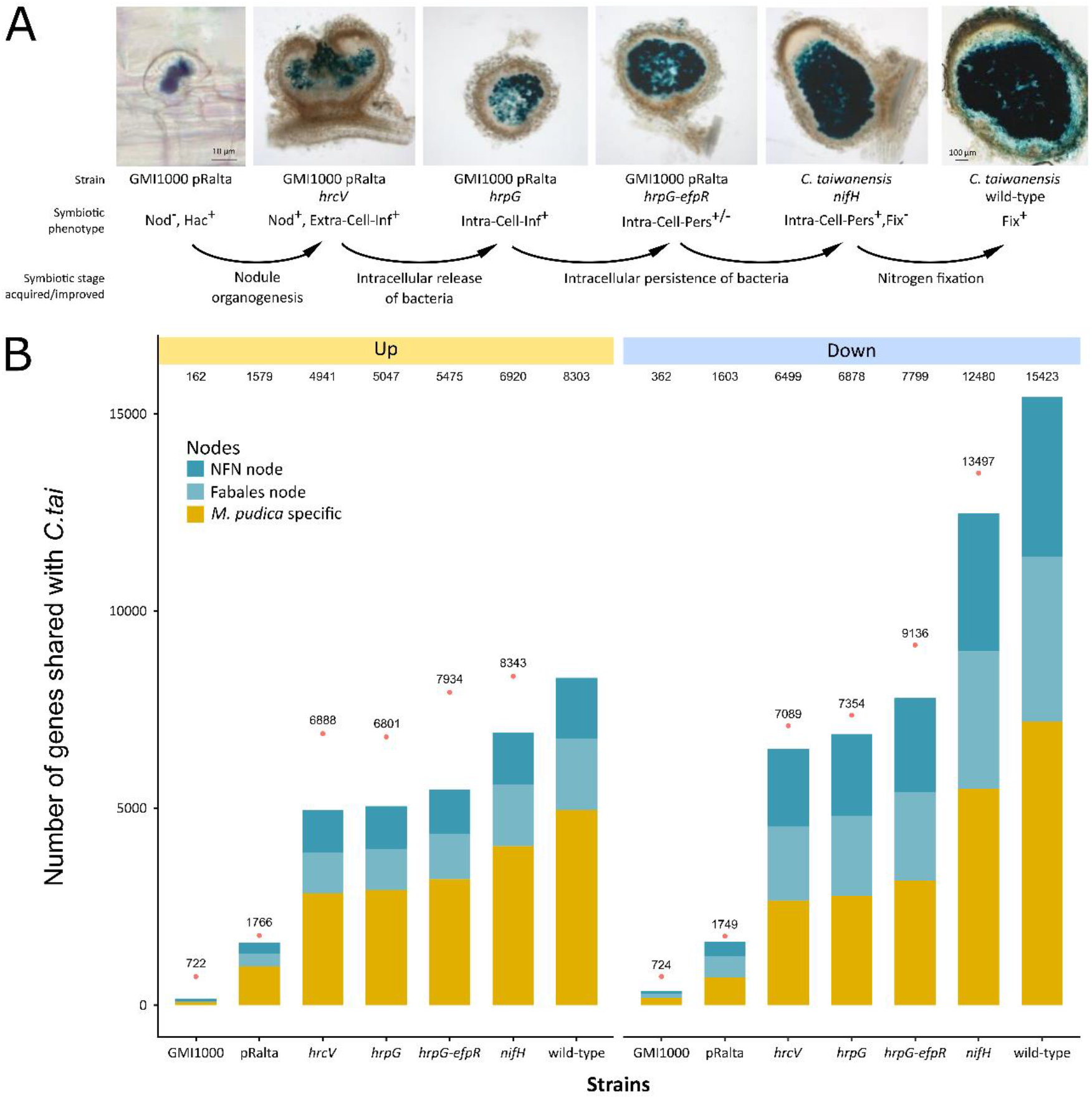
Genes recruited along the experimental evolution of RNS. (A) Symbiotic phenotypes of *Ralstonia* and *Cupriavidus* symbionts. *M. pudica* plants were inoculated with *lacZ*-tagged strains and nodules were harvested at 10 dpi for *Ralstonia* and *C. taiwanensis nifH* mutants and at 14 dpi for *C. taiwanensis* wild-type strain. Roots and nodule sections were stained with X-gal. The *C. taiwanensis* wild-type picture is from42. Nod, nodule formation. Hac, root hair curling. Extra-Cell-Inf, extracellular infection of nodules. Intra-Cell-Inf, intracellular infection of nodules. Intra-Cell-Pers, intracellular persistence. Fix, nitrogen fixation. (B) Number of genes that are up and down regulated in nodules formed by the different *Ralstonia* and *Cupriavidus* mutants and shared with the symbiotic response obtained with the *C. taiwanensis* wild-type strain. The distribution of these genes in the NFN and Fabales nodes and in the *M. pudica* specific gene set is indicated. Pink dots indicate the total number of DEG.

The evolution of improved symbiotic abilities in *Ralstonia* strains correlated with a gradual increase in the number of *M. pudica* DEG that are also DEG during the interaction with the wild-type *C. taiwanensis* strain (Figure 2B). The gain of the symbiotic plasmid was sufficient on its own to activate 19%/10% (up/down) of the whole symbiotic response (Figure 2B, Table S4). Accompanying this gain of symbiotic response, the GMI1000+pRalta strain also did not activate the expression of 553 *M. pudica* genes specifically induced by the wild-type GMI1000 *R. solanacearum* strain (Figure 2B). A significant number of these genes are associated with GO terms potentially linked to defense mechanisms such as “oxidation-reduction process”, “cell wall organization”, and “metabolism of secondary metabolites” (Table S7). This indicates that the horizontal gain of a symbiotic plasmid, a phenomenon widely observed within rhizobial populations^43^, may be sufficient to limit the activation of plant immunity. The *Ralstonia* nodulating but non-infective strain *hrcV* increased the shared response with the wild-type symbiont up to 60%/42% (up/down) while the well-infecting strain *hrpG-efpR* reached 66%/50% (up/down) of the wild-type symbiotic response. This pattern confirms phenotypic observations indicating that evolved *Ralstonia* strains are arrested at different stages along the progression towards a fully functional mutualistic state.

### The ancestral transcriptomic signature of symbiont perception, nodule organogenesis and N_2_-fixation

Next, we compared the plant responses to bacterial strains able/unable to reach different symbiotic traits (Figure 2A, Table S9) to identify *M. pudica* genes whose expression is associated with these different traits (response to NF, nodule organogenesis, intracellular release of bacteria (hereafter called “Release”), intracellular persistence (hereafter called “Persist”) and nitrogen fixation) (Figure 3A and Table S4). Transcriptomic responses to direct NF treatments were also available for two other Fabales, *Medicago truncatula* and *Lotus japonicus*^44,45^. Another dataset is available for *Medicago truncatula* obtained from laser-capture microdissections of the different nodule cell layers and corresponding to symbiosome release (FIId) and intracellular maintenance (FIIp), plant and bacteroid differentiation (FIIp, IZ) and nitrogen fixation (ZIII) (Table S3 and S4)^46^. To consider genes related to the different traits, genes have to be deregulated in the same way in the whole symbiotic transcriptomic response (Figure 3, Table S3 and S4), allowing us to position them in their evolutionary context.

**Figure 3:**
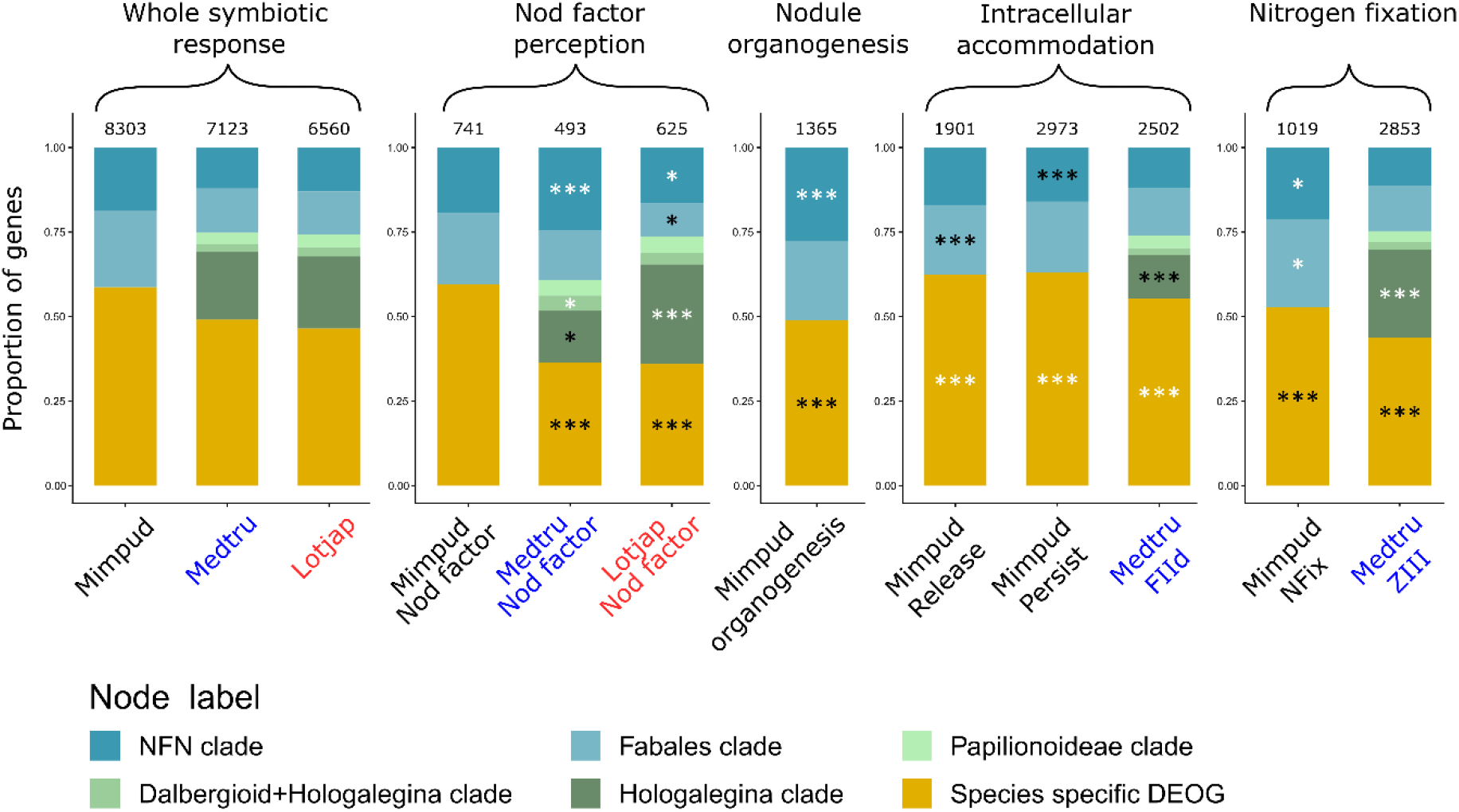
Evolutionary symbiotic stages response. Barplots representing the proportion of genes that are up regulated in whole symbiotic response and the different symbiotic traits, in the different evolutionary nodes. Mimpud, *Mimosa pudica*. Medtru, *Medicago truncatula*. Lotjap, *Lotus japonicus*. Asterisks indicate if the proportion is significantly different from the same node in whole symbiotic response using Fisher exact test, white for higher proportion and black for lower proportion. *p*-values: *0.05 > P > 0.01, **0.01 > P > 0.001, ***P < 0.001, absence of symbols: non-significant.

The distribution of the gene sets for the different traits in the different evolutionary nodes were compared to the whole symbiotic transcriptomic response (Figure 3). To do so, we used Fisher exact test between the whole symbiotic transcriptomic response and each trait, node by node (Figure 3 and Table S9), to estimate over/under representation of genes in the different nodes. This analysis indicates that the molecular basis of all stages of RNS involves genes that were already acting in the most recent common ancestor of all RNS-forming species, although in different proportions.

Perception of symbiont-produced Nod-factors in *M. truncatula* and *L. japonicus* as well as nodule organogenesis-associated genes were enriched in ancestral genes. In addition, both processes were linked with an impoverishment of species-specific DEOGs (Figure 3A and Table S9). Interestingly, 71 DEOGs shared by at least two species belong to the ancestral node (Figure S1), representing more than 50% of the shared DEOGs. Taken together, these results suggests that the Nod-factor perception recruited ancestral DEOGs followed by a large diversification to allow species specific refinement to facilitate early signaling and facilitate recognition between symbiotic partners. Among the ancestral DEOGs in response to Nod-factors, we detected the well-characterized NF-signaling components such as the transcription factors NIN, NF-YA1, NF-YA2 and ERN1/2, the infection genes RPG, VAPYRIN, SYFO and the LysM-RLK EPR3/LYK10^47^, the LRR-RLK RINRK1^48^ or the cytosolic kinase symCRK (Figure 4A, Table S4 and S7). The chitinase CHIT5 known to play a role in NF turn-over in the Fabales *L. japonicus*^49^ was also found as part of this shared NF-response, indicating that modulating NF levels was part of the ancestral RNS.

**Figure 4:**
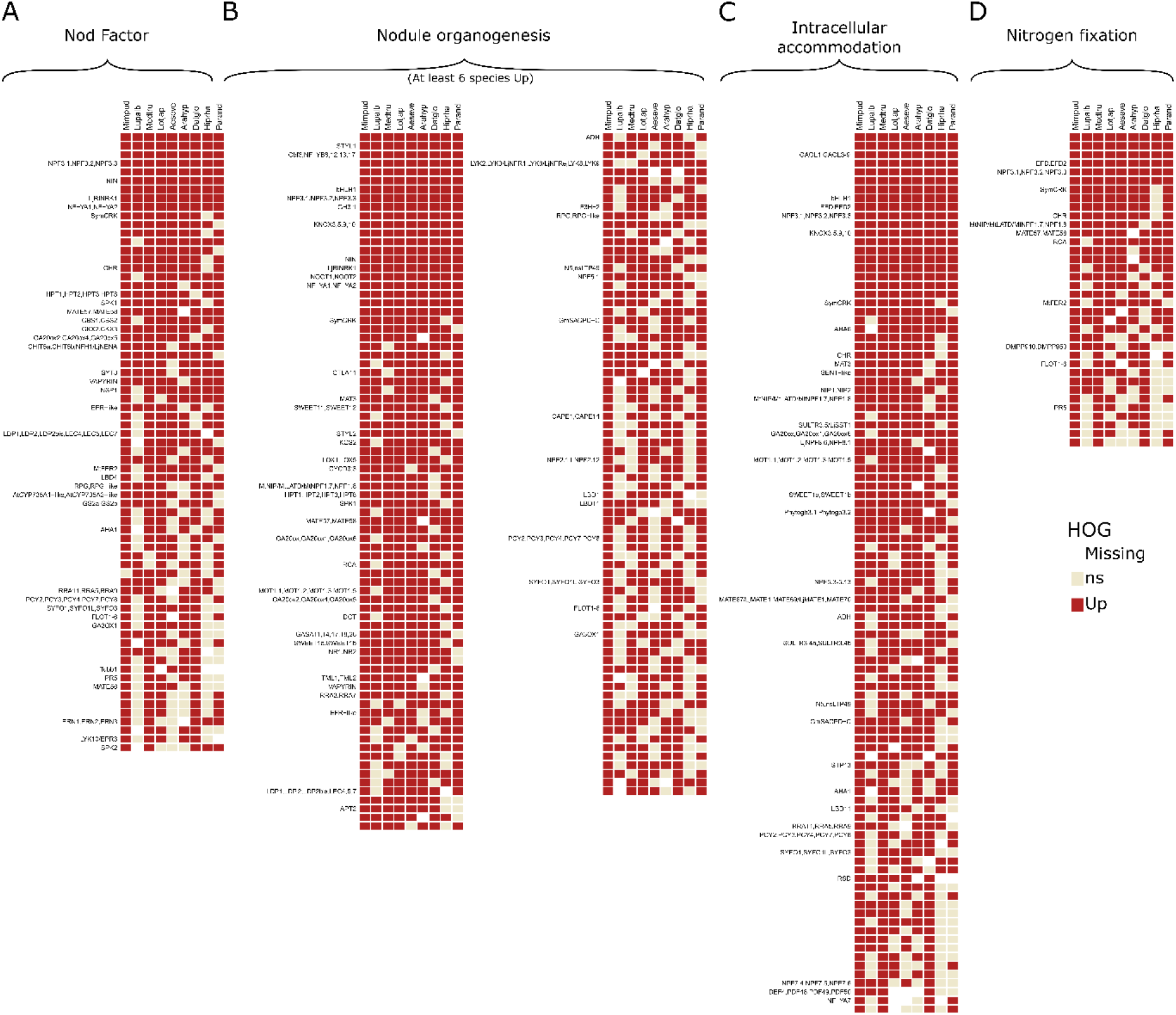
Heatmap representing the phylo-transcriptomic pattern for each for the 9 species concerning DEOGs assigned to NFN ancestral node. (A) Nod factor related DEOGs. (B) Nodule organogenesis related DEOGs. Only DEOGs shared by 6 species are displayed for graphical purpose. (C) Intracellular accommodation (including symbiosome release and intracellular persistence) related DEOGs. (D) Nitrogen fixation related DEOGs.

Organogenesis has been scrutinized in model legumes, revealing genes, in particular transcription factors, essential for the formation and maintenance of the nodule identity^50^. Many of these transcription factors were recovered in the inferred ancestral transcriptomic signature of organogenesis (Figure 4A, Table S4 and S7). Expectedly, genes involved in this module partially overlap with the NF-responsive genes, including the master regulator *NIN* and its direct or indirect targets *RPG*, *NF-YA1*, *NF-YA2* and *ERN1/2*, while other NIN targets, such as the transcription factors of the NF-YB family, LBD11 or STY1/2 involved in the production of auxin maxima required for nodule primordium emergence^51,52^, specifically belong to the organogenesis module (Figure 4B). Another well-known transcription factor, KNOX3, regulating nodule development through activation of cytokinin biosynthesis but acting upstream of NIN was found as part of this ancestral organogenesis program^53^. Finally, NOOT1 and NOOT2, which are known to maintain nodule identity in diverse legumes, were also detected^54^. Besides the known genes, 31 orthogroups annotated as transcription factors, so far not analyzed in the context of RNS, were detected. Their position in the organogenesis program remains to be determined (Figure 4B).

Nitrogen-fixation is a unifying feature of RNS. However, it has been predicted to be a trait that experienced significant refinement during the diversification of the NFN (Figure 4D). Mechanisms by which conditions for nitrogen fixation by the diverse symbionts (*Frankia*, alpha- and beta-proteobacteria) are provided in the nodules vary significantly. Despite this diversification, our analysis revealed an over-representation of DEOGs associated with N_2_-fixation and linked with the ancestral RNS gene set for *M. pudica* (Figure 3A) and less species specific DEOGs identified in both *M. pudica* and the nodule ZIII of *M. truncatula* (Figure 3 and Table S9). Most of these genes encode enzymes that have not been characterized yet (Table S4).

Although the mixed step of intracellular accommodation including the release of symbiosomes and their persistence in the *Medicago* and *Mimosa* dataset displays a peculiar evolutionary pattern (see below and Figure 3), this symbiotic stage also involved genes that are part of the ancestral transcriptomic response. Suppressors of plant defense in nodules, SYMCRK^55^ and RSD^56^, as well as the transcription regulator EFD required for both plant and bacteroid differentiation in *Medicago*^57,58^ participate to this ancestral response. Looking specifically at the *M. pudica* data, we found VAPYRIN, RPG, some flotillin and remorin genes and the syntaxin SYN, which are well-known infection-associated genes^33,59–61^. We thus hypothesize that a proportion of genes linked with the ancestral RNS transcriptome and associated with intracellular accommodation reflects infection (Figure 4C). DEOGs related to intracellular accommodation have to be identified in the “Release” and/or “Persist” DEOG set of *M. pudica* and “FIId” DEOG set of *M. truncatula* (Table S3 and S4).

As mentioned for the response to Nod-factors and organogenesis, DEOGs identified for the different traits often overlap suggesting that genes, such as transcription factors, may act at different symbiotic stages^50^.

We identified the gene modules associated with ancient symbiotic processes including genes whose position in the symbiotic pathway remains to be characterized. Altogether, this indicates that the mechanisms governing the response to Nod-factors, nodule organogenesis and nitrogen-fixation in extant RNS-forming species has been likely conserved since their most recent common ancestor

### Signature of convergent evolution for symbiosome formation in legumes

By contrast with symbiont perception, nodule organogenesis and nitrogen-fixation, the evolutionary pattern of the intracellular accommodation of rhizobia showed a decreased link with ancestral genes and an enrichment in species-specific DEOGs in both *M. pudica (*Mimosoideae) and *M. truncatula* (Papilionoideae, Figure 3A and Table S9). Compared to other orders of the NFN clade, RNS is evolutionary stable in the Fabales subfamillies Mimosoideae and Papilionoideae. It has been hypothesized that this stability may be linked with the occurrence of released symbiosomes exclusively found in these two clades^10^. Such a trait distribution might either reflect an ancestral gain in the Fabales and multiple subsequent losses, or be the result of convergent evolution. The fact that the transcriptomic signature associated to that stage depends much more on genes regulated in a species-specific manner in both *M. truncatula* and *M. pudica* than the other, ancestral, traits strongly supports the hypothesis of convergent gains of symbiosome formation in the two lineages (Figure 3A and 5). This species-specific transcriptomic change may be the results of either the recruitment of existing genes into the symbiotic transcriptomic response or the recruitment of genes that have evolved *de novo* in each lineage.

**Figure 5:**
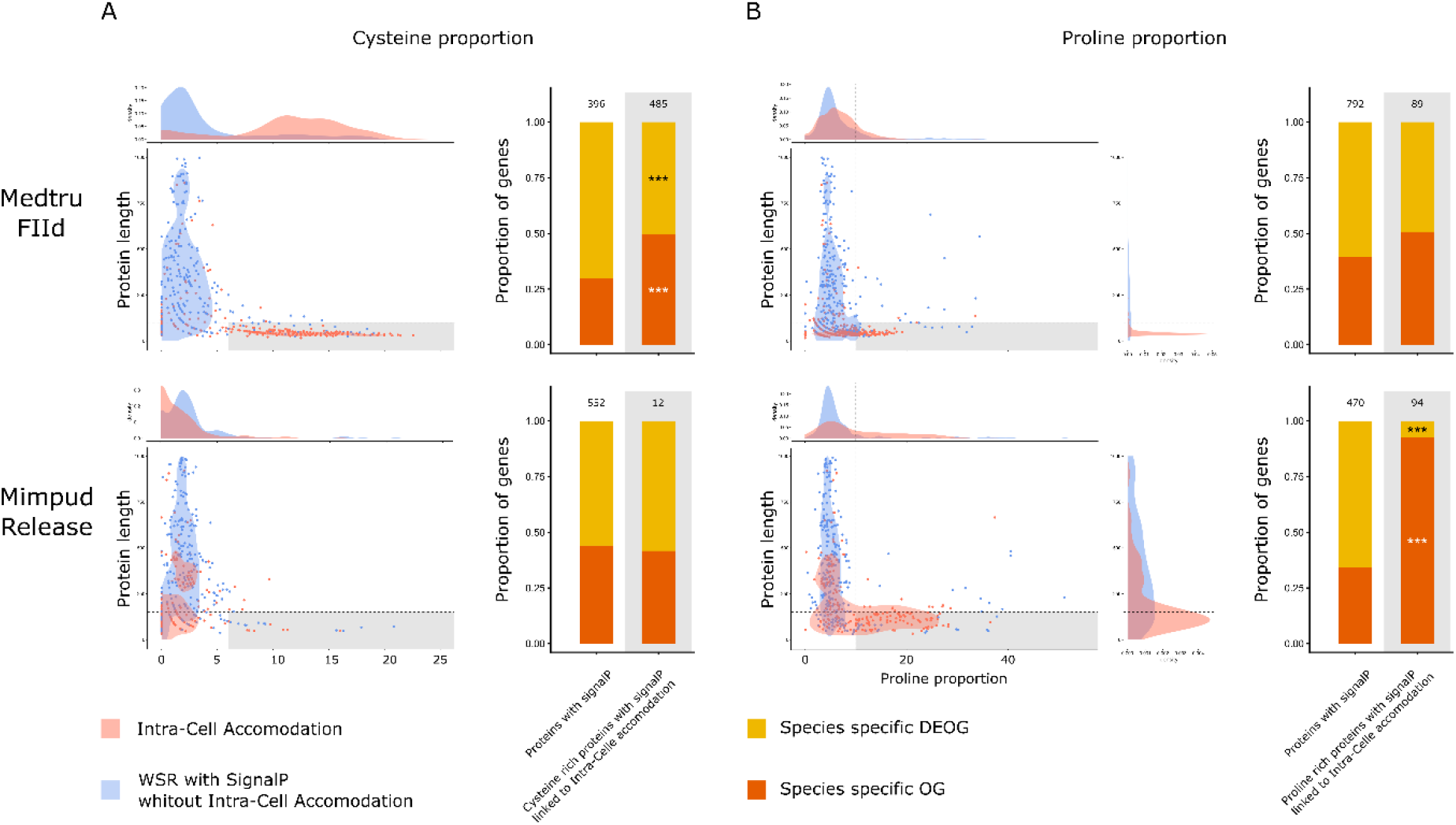
Characteristics of proteins with signal peptide from species-specific DEOGs related to intracellular accommodation in *M. truncatula* and *M. pudica*. (A) Scatterplot representing proportion of cysteine (x axis) and protein length (y axis) for up regulated genes in whole symbiotic response (WSR) (blue) and intracellular accommodation (red). Barplots representing the proportion of genes that are up regulated in whole symbiotic response without intracellular accommodation responsive genes in species-specific DEOGs and species-specific OGs. Asterisks indicate if the proportion is significantly different from the same node in whole symbiotic response using Fisher exact test, white for higher proportion and black for lower proportion. *p*-values: *0.05 > P > 0.01, **0.01 > P > 0.001, ***P < 0.001, absence of symbols: non-significant. (B) Scatterplot representing proportion of proline (x axis) and protein length (y axis) for up regulated genes in whole symbiotic response (blue) and intracellular accommodation (red). Barplots representing the proportion of genes that are up regulated in whole symbiotic response without intracellular accommodation responsive genes in species-specific DEOGs and species-specific OGs. Asterisks indicate if the proportion is significantly different from the same node in whole symbiotic response using Fisher exact test, white for higher proportion and black for lower proportion. *p*-values: *0.05 > P > 0.01, **0.01 > P > 0.001, ***P < 0.001, absence of symbols: non-significant.

We determined the nature of the genes associated to these species-specific responses. First, when compared to the whole symbiotic response, proteins related to intracellular accommodation (Release, Persist and FIId) and presenting a signal peptide are more abundant, in proportion, in both *M. pudica* and *M. truncatula* (Table S9). We looked into the length and amino acid composition of these proteins with signal peptide sequences. We observed significantly shorter proteins compared to the whole response with signal peptides proteins for both *M. pudica* (mean_Release_=204.1 vs mean_Mimpud_=404.6, Figure 5 and Table S9) and *M. truncatula* (mean_FIId_=66.7 vs mean_Medtru_=297.1, Figure 5 and Table S9). In addition, we found that these proteins showed more cysteine residues in *M. truncatula* and more proline residues in *M. pudica* (Figure 5 A and B, Table S9). Following these trends, we observed an enrichment in species-specific orthogroups (*i.e.* only the sequence of the given species is present in the orthogroup) for cysteine rich and small proteins in *M. truncatula*, but not for *M. pudica* (Figure 5A and Table S9). In *M. truncatula*, intracellular accommodation is partly mediated by small proteins, with a signal peptide and containing a high proportion of Cysteine known as Nodule Cysteine-Rich peptides (Figure 5A). In inverted repeat-lacking clade (IRLC) and some Dalbergioid legumes, these small secreted peptides have been shown to trigger the terminal differentiation of the nitrogen-fixing symbionts *via* antimicrobial activities preventing bacteroid proliferation outside the plant^62,63^. These NCRs corresponds to the species-specific genes identified here for *M. truncatula*. Reversely, we observed an enrichment in species-specific orthogroups for proline-rich and small proteins in *M. pudica*, but not for *M. truncatula* (Figure 5B and Table S9). Proline-rich peptides have been found in insects, mammals and plants where they play a role as antimicrobial compounds^64,65^. Although *M. pudica* symbionts are not terminally differentiated, their revivability outside the plant is limited^42^. The actual function of these proline-rich short proteins remains to be determined. Altogether, the presented data support that the convergence in the intracellular accommodation of symbionts evolved by, at least, two independent but analogous molecular processes, the *de novo* evolution of nodule-induced small proteins already proposed in Dalbergioid and IRLC^63^.

## Conclusion

From the distribution of the trait and phylogenomic analyses, the leading hypothesis for the origin of RNS is that it was gained before the radiation of the NFN clade approximately 110 million years ago^13,14^. Here, we propose that RNS in that most recent common ancestor looked very similar to RNS in extant species. With the ancestral RNS transcriptomic signature now defined, future studies will have to decipher how this state evolved from a non-RNS-forming state. A role for the common symbiosis pathway in that process can be anticipated given its phylogenetic link with all the plant intracellular symbioses^66^, and the reverse genetic data obtained in diverse RNS-forming species^9^. The gain of a regulatory link between the common symbiosis pathway and the central RNS-regulator NIN at the base of the NFN clade represents one of the events, which have played a role in the transition from the non-NFN-forming to the NFN-forming state^67^. For millions of years, RNS has been maintained in diverse lineages of the Fagales, Fabales, Cucurbitales and Rosales with presumably very high rates of symbiosis loss. In only two lineages, the sub-families Mimosoideae and Papillionideae in the Fabales order, RNS has become evolutionary stable^10^. Our results support the hypothesis that evolutionary stability was acquired through the convergent evolution of symbiosome release and enhanced control of the bacterial symbiont gained by the expansion of putative antimicrobial peptide gene families. Besides reconstructing the ancestral RNS state, our comparative transcriptomic approach has determined the blueprint for RNS that can be considered as the target to be reached by the multiple consortia ambitioning to engineer nitrogen-fixing symbiosis in crops^68–70^.

## Methods

### *Mimosa pudica* genome assembly and annotation

#### High Molecular Weigh DNA extraction

DNA was isolated from frozen young leaves using QIAGEN Genomic-tips 100/G kit (Cat No./ID: 10243). We have followed the tissue protocol extraction. Briefly, 1 g of young leaf material were ground in liquid nitrogen with mortar and pestle. After 3 h of lysis at 50 °C with proteinase K and one centrifugation step, the DNA was immobilized on the column. After several washing steps, DNA is eluted from the column, then desalted and concentrated by alcohol precipitation. The DNA is resuspended in TE buffer.

### *Mimosa pudica* whole genome sequencing: PacBio library preparation

On HMW DNA sample, a standard SMRTbell® library was constructed using the SMRTbell Template Prep kit 1.0 (Pacific Biosciences, Menlo Park, CA, USA) according to PacBio recommendations PN 100-938-400-03).

HMW DNA was sheared using Megaruptor 2 system (Diagenode, Liège Science Park, Belgium) to obtain a 40 Kb average size. Following an enzymatic treatment on 7.5 μg of sheared DNA sample for DNA damage repair, ligation with hairpin adapters to both ends of the targeted double-stranded DNA (dsDNA) molecule was performed to create a closed, single-stranded circular DNA. A nuclease treatment was performed by using SMRTbell Enzyme Clean-up kit (Pacific Biosciences, Menlo Park, CA, USA). A size-selection with Blue-Pippin system (Sage Science, Beverly, MA, USA) to remove fragments less than 15 Kb was done on purified sample with 0.45× AMPure PB beads (Pacific Biosciences, Menlo Park, CA, USA). The size and concentration of the final library were assessed using the Fragment Analyzer system (Agilent, Santa Clara, CA, USA) and the Qubit Fluorometer and Qubit dsDNA HS reagents Assay kit (Thermo Fisher Scientific, Waltham, MA, USA), respectively.

Sequencing primer v3 and Sequel DNA Polymerase 3.0 were annealed and bound, respectively to the SMRTbell library. The library was loaded on 8 SMRTcells 1M and sequencing was performed on the Sequel I system with Sequel Sequencing kit 3.0, a run movie time of 600 min and Software v6.0 (PacBio).

### Genome assembly

The genome was assembled in three steps.

In Step 1, Canu version 1.8^71^ was used to trim, correct and assemble the 5 815 198 subreads for a total size of 85 Gb, i.e. an estimated coverage of 94×. Program parameters were corOutCoverage=40, minReadLength=1000 and an input genome size estimate of 900Mb.

In Step 2, the raw data, from PacBio Sequel bam files were aligned on the draft assembly. For this step we used the wrapper of minimap2^72^, pbmm2 include in the SMRT Analysis Software version 7.0.0 (https://www.pacb.com/products-and-services/analytical-software/smrt-analysis/). Program parameters were « pbmm2 align --preset ‘SUBREAD’ -c 70 -l 500 ».

In the final step (Step 3), we used this mapping result in order to polish the draft assembly and generate a high-quality final assembly. For this step we used the variantCaller command, include in the SMRT Analysis Software version 7.0.0, with the arrow algorithm. Program parameters were --algorithm arrow --minConfidence 40 --minCoverage 70 --coverage 100 -- minReadScore 0.65.

The final polished assembly produced have a total of 1343 contigs with a total length of 842 189 795 pb, a largest contig of 24 170 175 pb and a N50 contigs of 13 257 064 pb.

### Preparation of ultra-high molecular weight (uHMW) DNA for Bionano optical mapping

Nuclei were purified from 0.5 g of dark treated young leaves according to the Bionano Plant tissue DNA isolation base protocol (# 30068 - Bionano Genomics) followed by uHMW DNA extraction based on the Bionano Prep SP kit (# 80030 - Bionano Genomics) and adapted by our laboratory for plant samples.

Briefly, plant leaves were flash frozen in liquid nitrogen and disrupted with a rotor-stator homogenizer (Qiagen). Nuclei were pelleted, washed and digested with proteinase K in lysis buffer. After the PMSF treatment, a centrifugation is added to eliminate the cell wall debris. The supernatant is precipitated with isopropanol and captured with magnetic disk (Nanobind disk). After several washes the uHMW DNA is eluted in elution buffer.

Labeling and staining of the uHMW DNA were performed according to the Bionano prep direct label and stain (DLS) protocol (30206 - Bionano Genomics). Briefly, labeling was performed by incubating 750 ng genomic DNA with DLE-1 enzyme (Bionano Genomics) for 2 hours in the presence of DL-Green dye (Bionano Genomics). The DLE 1 enzyme recognizes the motif CTTAAG. Following proteinase K (Qiagen) digestion and non-fixed dye cleanup by membrane adsorption, the DNA backbone was stained with DNA Stain solution (Bionano Genomics) and incubated overnight at room temperature. The labelled DNA concentration was measured with the Qubit dsDNA HS assay (Invitrogen).

### Data collection, optical map construction and genome scaffolding

Labelled DNA was loaded on the Saphyr G1 chip according to Saphyr System User Guide (30247 - Bionano Genomics). Data processing was performed using the Bionano Genomics Access software https://bionanogenomics.com/support-page/bionano-access-software/).

480 Gb of molecules larger than 150 Kb with a N50 of 199 kbp were produced and represented 533×. It corresponded to 533× coverage of the 900 Mb estimated size of M. pudica genome. These molecules were assembled using RefAligner with default parameters. It produced 110 genome maps with a N50 of 16.1 Mbp for a total genome map length of 833 Mbp.

Finally, a hybrid scaffolding was performed between the polished PacBio assembly and the optical genome maps with hybridScaffold pipeline with default parameters. We obtained 74 hybrid scaffolds ranging from 128 kbp to 25.5 Mbp (total length 797 Mbp with N50 = 16.1 Mbp).

### *Mimosa pudica* genome structural annotation

The *Mimosa pudica* gene models were predicted by the eukaryotic genome annotation pipeline egn-ep (http://eugene.toulouse.inra.fr/Downloads/egnep-Linux-x86_64.1.5.1.tar.gz) using trained statistical models adapted for plants (http://eugene.toulouse.inra.fr/Downloads/WAM_plant.20180615.tar.gz). This pipeline manages automatically probabilistic sequence model training, genome masking, transcript and protein alignments computation, alternative splice sites detection and integrative gene modelling by the EuGene software release 4.2a ^73^(http://eugene.toulouse.inra.fr/Downloads/eugene-4.2a.tar.gz).

Four protein databases were used to detect translated regions: i) the proteome of *Medicago truncatula* A17 version 5 annotation release 1.6 https://medicago.toulouse.inra.fr/MtrunA17r5.0-ANR/), ii) the proteome of the previous *Mimosa pudica* Illumina genome^14^, iii) Swiss-Prot - October 2016 and iv) the proteome of *Arabidopsis thaliana* TAIR10 version. Proteins similar to REPBASE were removed from the three datasets (to avoid the integration of TE related proteins in the training steps). Chained alignments spanning less than 50% of the length of the database protein were removed. The proteome of *M. truncatula* release 1.6 was used as a training proteome by EuGene.

Three input transcripts for EuGene were used. One transcriptome based on mapping predicted from the 136 RNAseq samples generated in this study (Table S1).

To obtain these transcripts, the raw fastq paired-end reads were cleaned by removing the adapters and the low-quality sequences using cutadapt^74^ (v2.1) and TrimGalore (v0.6.5, https://github.com/FelixKrueger/TrimGalore) with -q 30 --length 20 options. The cleaned reads were mapped against the Mimosa pudica genome assembly using HISAT2^75^ (v2.1.0) with -- score-min L,−0.6,−0.6 --max-intronlen 10000 --dta --rna-strandness RF options. Duplicated reads were removed using SAMtools^76,77^ (v1.9) markdup command. Transcripts were predicted using Stringtie ^78^ (v2.1.4) with --fr -f 0.8 on each sample. All 80 gtf sample files were merged together using stringtie --merge with standard options. Transcripts fasta file was generated using gffread^78,79^ (v0.11.6) with -w option.

We also *de novo* predicted two transcriptomes from two batches of 10 samples of our same RNAseq data using DRAP pipeline^80^ (v1.92, http://www.sigenae.org/drap). runDrap was used on the 20 samples applying the Oases RNAseq assembly software^81^. runMeta was used to merge assemblies without redundancy based on predicted transcripts with fpkm 1. These transcriptomes were employed as a training transcriptome by EuGene. Finally, 73,541 protein-coding genes, 1107 tRNAs, 114 rRNAs and 3913 ncRNAs were annotated.

Genome assembly, annotation file and gene models are publicly available through MyGenomeBrowser^82^ https://bbric-pipelines.toulouse.inra.fr/myGenomeBrowser?share=manage&filter=Mimpud_MpudA1P6v1 and through NCBI under BioProject PRJNA787464.

### RNAseq data

To decipher the ancestral transcriptomic response of root nodule symbiosis species we generated new RNAseq data for *Mimosa pudica* and *Lupinus albus*.

### *Mimosa pudica* RNA isolation and sequencing

*Mimosa pudica* tissue samples were harvested at 1, 3 and 5 days post-inoculation (dpi) for non-inoculated plants, at 1 and 3 dpi for plants inoculated with non-nodulating *R. solanacearum* strains, at 1, 3, 5, 7 and 10 dpi for plants inoculated with *R. solanacearum* strains and at 1, 3, 5, 7, 14 and 21 dpi for plants inoculated with *C. taiwanensis* strains (Table S1). Samples from four independent biological replicates were harvested at each time point. Total RNA was isolated from roots, nodule primordia, and nodules using a NucleoSpin RNA Plus kit (Macherey-Nagel) according to the manufacturer’s instructions, treated with a rDNase (Macherey-Nagel) for 10 min at 37°C and then cleaned up with the NucleoSpin RNA clean-up kit (Macherey-Nagel). RNA quality was verified on a 2100 Bioanalyzer Instrument (Agilent) and quantified on a QubitTM fluorometer (Thermo Fisher Scientific). RNA sequencing was performed at the GeT-PlaGe core facility, INRAE Toulouse. Polyadenylated mRNA and RNA-seq libraries have been prepared according to Illumina’s protocols using the Illumina TruSeq Stranded mRNA sample prep kit to analyze mRNA. Briefly, mRNA were selected using poly-T beads. Then, RNA were fragmented to generate double stranded cDNA and adaptators were ligated to be sequenced. 11 cycles of PCR were applied to amplify libraries. Library quality was assessed using a Fragment Analyser and libraries were quantified by QPCR using the Kapa Library Quantification Kit. RNA-seq experiments have been performed on an Illumina NovaSeq 6000 using a paired-end read length of 2×150 pb with the Illumina NovaSeq 6000 sequencing kits.

### *Lupinus albus* RNA isolation and sequencing

*Lupinus albus* tissue samples were harvest at 21dpi for non-inoculated plants and inoculated plants with Bradyrhizobium sp. 1AE200 strain. Three biological replicates of inoculated and non-inoculated *Lupinus* root samples were used for RNA sequencing. Samples were grounded using pestle and mortar. RNA extraction and DNase treatment were performed using respectively E.Z.N.A RNA extraction kit (Omega-Biotek) and TURBO DNA kit (Invitrogen) according to the manufacturer’s instructions. Quality of RNAs was assessed using the Agilent 2100 Bioanalyzer system. RNA sequencing was performed by the Eurofins genomics facility. Polyadenylated mRNA and RNA-seq libraries have been prepared according to Illumina’s protocols using the Illumina TruSeq Stranded mRNA sample prep kit to analyze mRNA. RNA-seq experiments have been performed on an Illumina NovaSeq 6000 using a paired-end read length of 2×150 pb with the Illumina NovaSeq 6000 sequencing kits.

### Differential gene expression analysis

All RNAseq libraries were mapped against their representative genome (Table S1) using nextflow^83^ (v20.11.0-edge) run nf-core/rnaseq^84^ (v3.0, 10.5281/zenodo.1400710) using *-profile debug,genotoul --skip_qc --aligner star_salmon* options. The workflow used bedtools^85^ (v2.29.2), bioconductor-summarizedexperiment (v1.20.0), bioconductor-tximeta (v1.8.0), gffread^79^ (v0.12.1), picard (v2.23.9), salmon^86^ (v1.4.0), samtools^76^ (v1.10), star^87^ (v2.6.1d), stringtie^78^ (v2.1.4), Trimgalore (v0.6.6) and ucsc (v377). Differencially expressed genes (DEGs) for the different species and experiments were estimated using ‘*edgeR*’^88^ in R^89^ (v4.1.2). Template script to estimate and identify DEGs is store on the following dedicated GitHub repository https://github.com/CyrilLibourel/Universal_nodulation_transcriptomic_response. Briefly, low expressed genes with less than 10 reads across each class of samples. After, gene counts were normalized by library size and trimmed mean of M-values (i.e. TMM) normalization method90. We estimated differentially expressed genes (DEGs) by comparing symbiotic states to non-inoculated roots for the different species. *Mimosa pudica* symbiotic traits (NF response, nodule organogenesis, bacterial release, intracellular persistence and nitrogen fixation) DEGs were analysed with the DicoExpress tool^91^, that relies on the R packages ‘*FactoMineR*’^92^ and ‘*edgeR*’^88^ to identify the genes that are differentially expressed between experimental conditions using generalized linear models. The lists of DEGs responsive genes as well as genes associated with the different symbiotic traits were determined by combining lists of DEG originating from multiple comparisons between 2 samples as indicated in the Table S3. Genes were considered differentially expressed when the FDR was below 0.05 (Benjamini-Hochberg correction) and specific log fold-change (see Table S3).

### Orthogroups reconstruction

To cross expression data among species we reconstructed orthogroups including the nine species for which RNASeq data during symbiosis are available as well as 14 representative species of each NFN clades and *Arabidopsis thaliana* as outgroup (Table S5). Orthogroups reconstruction was performed with OrthoFinder v2.5.2 (10.1186/s13059-019-1832-y) with the ultra-sensitive Diamond mode (-S diamond_ultra_sens option). The inferred tree by OrthoFinder was then checked for consistent reconstruction with known species tree and OrthoFinder reran based on the species tree using alignment and phylogeny method with mafft v7.313 (10.1093/molbev/mst010) and fasttree v2.1.10 (10.1093/molbev/msp077) respectively to infer orthogroups.

### Statistical analysis

The different scripts used to cross orthogroups and DEGs to estimate differentially expressed orthogroups (DEOGs), reconstruct the ancestral transcriptomic symbiotic response, identify symbiosome accommodation proteins and all statistical related analysis are freely available on the following dedicated GitHub repository https://github.com/CyrilLibourel/Universal_nodulation_transcriptomic_response.

In order to estimate the null distribution of the number of DEOGs for each node, we first determined to which evolutionary node each orthogroup belongs using a maximum parsimony ancestral state reconstruction for discrete traits^24^ (i.e. presence/absence of species genes in orthogroup). For each species, we randomly select genes accordingly to (i) the number of DEGs estimated in the species and (ii) the node that belongs the orthogroup containing DEGs. We did this process for the nine species and determined to which evolutionary node each orthogroup belongs using the same maximum parsimony ancestral state reconstruction (i.e. presence/absence of species DEGs in orthogroup). We did this process 1000 times to get the null expectation (Table S6).

## Data and code availability

Scripts used to cross orthogroups and RNAseq data are available on the GitHub page https://github.com/CyrilLibourel/Universal_nodulation_transcriptomic_response.

*Mimosa pudica* genome assembly and raw sequenced data are available on NCBI under BioProject PRJNA787464.

All the repositories of RNAseq used in this study are detailed in the Table S1.

## Acknowledgements

The authors thank Oswaldo Valdes-Lopez, Pascal Gamas and the ENSA consortium for discussions and feedback on the manuscript, and B. Perret for the *Lupinus albus* seeds. We are grateful to the genotoul bioinformatics platform Toulouse Occitanie (Bioinfo Genotoul, https://doi.org/10.15454/1.5572369328961167E12) for providing computing resources. This study was supported by the Fédération de Recherche Agrobiosciences, Interactions et Biodiversité, the French National Research Institute for Agriculture, Food and Environment (INRAE, Plant Health and Environment Division), the “Laboratoires d’Excellence (LABEX)” TULIP (ANR-10-LABX-41)” and by the “École Universitaire de Recherche (EUR)‘ TULIP-GS (ANR-18-EURE-0019). J.K., C.L. and P-M.D are supported by the project Engineering Nitrogen Symbiosis for Africa (ENSA) currently funded through a grant to the University of Cambridge by the Bill & Melinda Gates Foundation (OPP1172165) and the UK Foreign, Commonwealth and Development Office as Engineering Nitrogen Symbiosis for Africa (OPP1172165). This project has received funding from the European Research Council (ERC) under the European Union’s Horizon 2020 research and innovation program (grant agreement No 101001675- ORIGINS) to P-M.D. PR has received funding from the EU in the framework of the Marie-Curie FP7 COFUND People Program, through the award of an AgreenSkills+ fellowship (under grant agreement n°609398) and from the European Union’s Horizon 2020 research and innovation program under the Marie Skłodowska-Curie grant agreement N° 845838.

## Author Contributions

CL, JK, PR, P-MD and DC designed the project. CL, JK, LK, A-CC, SC, TV, JMC, CC, ID, SC, WM, TB, AS, CM-B, PR, P-MD and DC conducted experiments. CL, JK, SC, PR, P-MD and DC analyzed data. CL, JK, PR, P-MD and DC wrote the manuscript with input from all authors.

## Supplementary Figures and Tables

**Figure S1:**
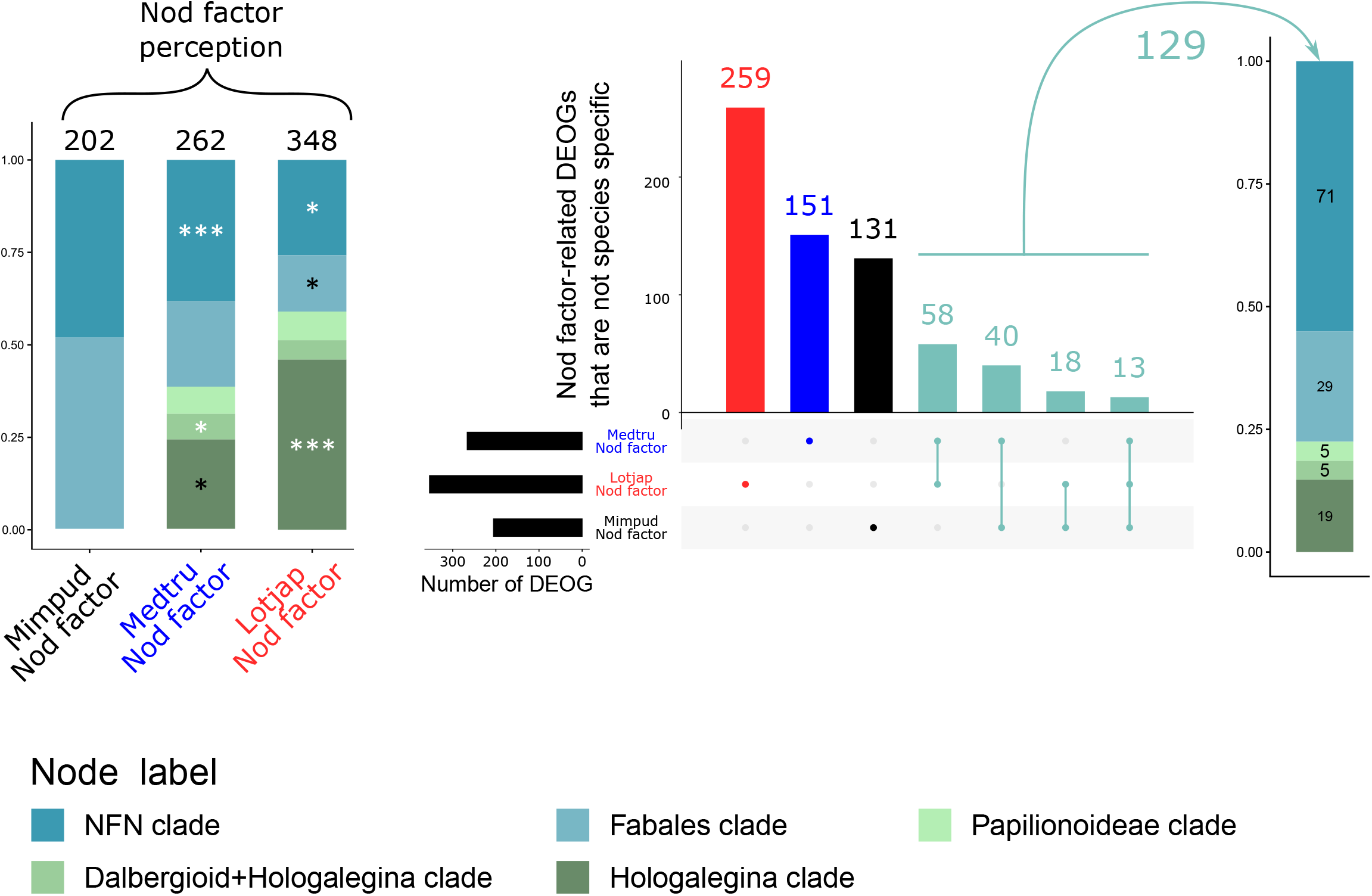
Shared ancestral Nod-factor response. (A) Barplots representing the proportion of genes that are up regulated in whole symbiotic response related to Nod factor perception, in the different evolutionary nodes, except for species specific DEOGs. Asterisks indicate if the proportion is significantly different from the same node in the whole symbiotic response using Fisher exact test, white for higher proportion and black for lower proportion. p-values: *0.05 > P > 0.01, **0.01 > P > 0.001, ***P < 0.001, absence of symbols: non-significant. (B) Upset plot representing overlap of DEOGs for L. japonicus, M. truncatula and M. pudica NF responsive orthogroups. Input lists correspond to DEOGs from panel (A) “Nod factor perception” without Species specific DEOGs. DEOGs specific to L. japonicus, M. truncatula and M. pudica are displayed in red, blue and black, respectively. Light blue: DEOGs shared by at least two species. (C) Proportion of shared DEOGs from panel (B) in the different nodes.

**Table S1: RNAseq samples used in this study.**

**Table S2: Raw sequence data and statistics for Mimosa pudica genome and annotation completeness**

**Table S3: Sample comparisons to identify DEGs associated with the different symbiotic traits or during the symbiotic interaction in the different plant species**

**Table S4: DEGs by species**

**Table S5: Species used for orthogroups reconstruction**

**Table S6: Observed and random expectation of DEOGs for each node**

**Table S7: List of orthogroups and their evolutionary position where the DEOG is assign**

**Table S8: GO terms enriched in the GMI1000-induced specific response**

**Table S9: Summary statistics of t-tests and Fisher tests**

**Table S10: Medicago truncatula and Mimosa pudica specific small secreted proteins associated with intracellular accommodation**

