## Supplementary material for "Comparative phylotranscriptomics reveals a 110 million years-old symbiotic program": Figure S1

Figure S1: Shared ancestral Nod-factor response. (A) Barplots representing the proportion of genes that are up regulated in whole symbiotic response related to Nod factor perception, in the different evolutionary nodes, except for species specific DEOGs. Asterisks indicate if the proportion is significantly different from the same node in the whole symbiotic response using Fisher exact test, white for higher proportion and black for lower proportion. p-values: *0.05 > P > 0.01, **0.01 > P > 0.001, ***P < 0.001, absence of symbols: non-significant. (B) Upset plot representing overlap of DEOGs for L. japonicus, M. truncatula and M. pudica NF responsive orthogroups. Input lists correspond to DEOGs from panel (A) “Nod factor perception” without Species specific DEOGs. DEOGs specific to L. japonicus, M. truncatula and M. pudica are displayed in red, blue and black, respectively. Light blue: DEOGs shared by at least two species. (C) Proportion of shared DEOGs from panel (B) in the different nodes.

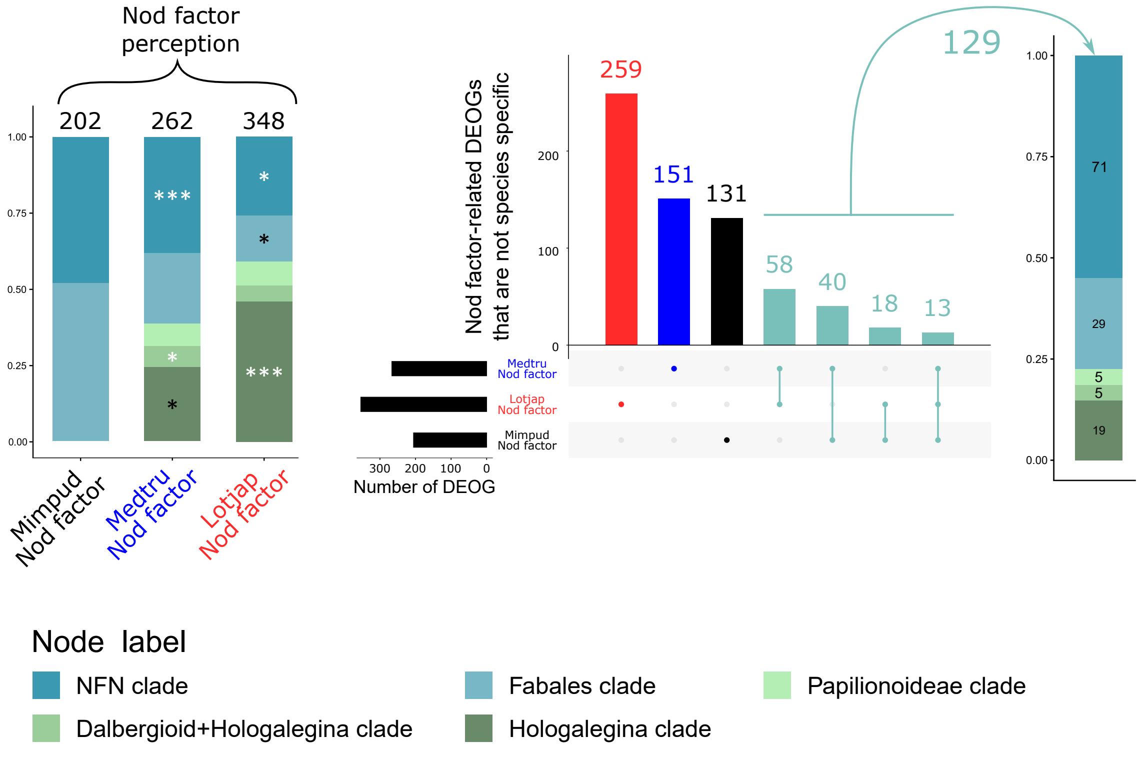
